# The Heterogeneous Nature of Atrioventricular Conduction Tissues in Tetralogy of Fallot demonstrated by Hierarchical Phase-contrast Tomography – redefining the anatomic substrate

**DOI:** 10.64898/2025.12.04.692356

**Authors:** V Sabarigirivasan, J Brunet, H Dejea, A Crucean, A Jegatheeswaran, C Capelli, E Pajaziti, Theresa Urban, Joanna Purzycka, Paul Tafforeau, C L Walsh, P D Lee, A C Cook

## Abstract

**Objectives:** Postoperative arrhythmias are frequent after Tetralogy of Fallot (ToF) repair, yet anatomic substrate and preventive strategies remain poorly defined. Using hierarchical phase-contrast tomography (HiP-CT) the atrioventricular conduction system in pediatric ToF specimens was investigated non-destructively and in 3D.

**Methods:** Eighteen whole-heart specimens (11 ToF, 7 controls) were imaged at the European Synchrotron, ESRF. Segmentation and 3D renderings demonstrated gross morphology. Morphology, size, depth and course of the non-branching bundle and right bundle branch (RBB) were quantified using custom computational pipelines. Segmentations were visualized in VheaRts, a Unity3D-based XR platform.

**Results:** The ToF conduction system was more draped than in controls, bilaterally spanning the septum, resembling early embryonic architecture. The RBB was variable in origin, course and morphologic structure with significantly smaller indexed cross-sectional area in native ToF versus controls (0.05 ± 0.20 vs 0.50 ± 0.20 mm², p = 0.005). One anomalous fasciculo-ventricular connection and six dead-end tracts were found. Regions at surgical risk included the posterior-inferior margin of the ventricular septal defect (VSD) and the septal crest along its nadir. The superior margin of the VSD at its intersection with the aortic root was free of conduction tissue. The HiP-CT to VR pipeline enabled interactive 3D visualization of conduction pathways relative to key structures.

**Conclusions:** This first pediatric cardiac HiP-CT series reveals a broader anomalous conduction complex in ToF, including variable RBB origin and hypoplasia, providing insight into preoperative vulnerability, arrhythmia, and surgical risk.

Figure 7:
Graphical Abstract

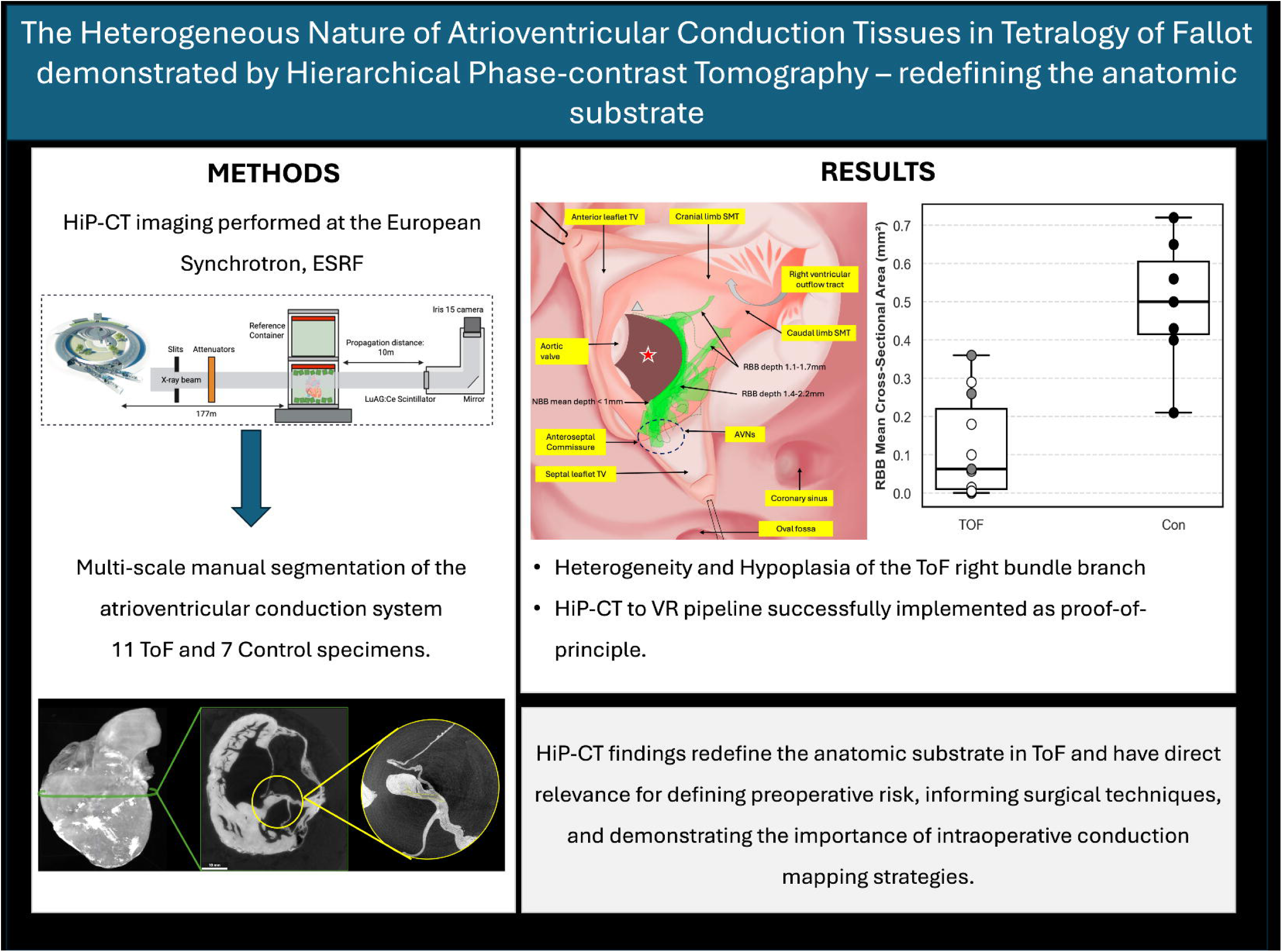

## Introduction

Postoperative arrhythmias represent a significant complication in Tetralogy of Fallot (ToF), with reported incidences reaching 43% in some multi-center studies^1,2^. Ventricular arrhythmias and conduction delays influence transient and long-term morbidity and mortality, but their etiology remains incompletely understood. Postoperative arrhythmias may result from arrhythmogenic changes in the atrioventricular conduction system (AVCS) or its surrounding myocardium causing junctional ectopic tachycardia (JET), or from direct injury during VSD closure or outflow tract resection^1,2^. Preoperative right bundle branch (RBB) block is rare (∼4%) but occurs in up to 60% of patients at discharge postoperatively. Prevalence increases to above 80% in childhood and adolesence^3^.

Current knowledge of the ToF AVCS system is based on seminal work in the 1950s-1980s which used destructive dissection and histologic serial sectioning of the ‘conduction block’ in small numbers of pediatric heart specimens^4–12^. While this laid the basis for surgical intervention, no additional data have been generated due to the lack of novel, non-destructive techniques. Hierarchical Phase-Contrast Tomography (HiP-CT) can image intact human organs with exceptional resolution using the phase-contrast capabilities of the BM18 beamline at the European Synchrotron, ESRF Extremely Brilliant Source^13^. It allows the non-destructive examination of whole organs, with local zooms down to histologic level (2μm voxel size), eliminating the need for physical subsampling^13^. Studies using other synchrotron imaging techniques or protocols have visualized the AVCS in whole fetal and pediatric specimens^14–16^, but only HiP-CT has been able to achieve this in intact adult hearts^17^. These investigations have been restricted to at most two specimens per study. We present the first whole-organ pediatric cardiac disease case series using HiP-CT. An overview of the study design is shown in the graphical abstract.

## Materials and Methods

### Specimen Description

Five hundred and fifty ToF hearts from the UCL Cardiac Archive Biobank (UK Human Tissue Authority Research License 12220) were reviewed, and eleven selected for imaging (Table 1). Hearts were chosen if they were within specific size constraints (105 mm × 105 mm × 167 mm), were in good condition on visual inspection, and possessed intact right ventricular outflow tracts (RVOT) and VSDs. Seven control specimens from Birmingham Children’s Hospital (BCH) (ethical approval: REC 22/PR/0906) & UCL were selected using the same criteria as above. All specimens were assigned a unique ID number under UCL CAB and transported to ESRF under transport authorization N° IE-2023-2966 issued by the French Ministry of Higher Education and Research.

**Table 1:** Morphologic characteristics of specimens included in this study.

| ID | Short-form link | Specimen Morphology |
| --- | --- | --- |
| UCL-ZCR-125 | T1 | PM outlet VSD, pulmonary stenosis |
| UCL-ZCR-173 | T2 | PM outlet VSD, absent pulmonary valve |
| UCL-ZCR-185 | T3 | PM outlet VSD, pulmonary atresia |
| UCL-ZCR-450 | T4 | PM and juxta-arterial VSD, pulmonary stenosis |
| UCL-ZCR-727 | T5 | PM outlet VSD, pulmonary stenosis, VSD patch + pericardial transannular patch |
| UCL-ZCR-769 | T6 | PM outlet VSD, absent pulmonary valve |
| UCL-ZCR-1770 | T7 | PM outlet VSD, pulmonary stenosis |
| UCL-ZCR-1771 | T8 | PM outlet VSD, pulmonary stenosis |
| UCL-ZCR-1840 | T9 | PM outlet VSD, pulmonary stenosis, VSD patch + dacron transannular patch |
| UCL-ZCR-2010 | T10 | PM outlet VSD, pulmonary stenosis, VSD patch + dacron transannular patch |
| UCL-ZCR-2577 | T11 | Muscular Outlet VSD, pulmonary atresia |
| UCL-ZCR-2359 | C1 | Normal |
| UCL-ZCR-2337 | C2 | Normal |
| UCL-ZCR-2625 | C3 | Normal |
| UCL-ZCR-2627 | C4 | Normal |
| UCL-ZCR-2628 | C5 | Normal |
| UCL-ZCR-2628 | C6 | Normal |
| UCL-ZCR-2630 | C7 | Normal |

### Sample Preparation and Data acquisition

HiP-CT imaging was performed at beamline BM18 (European Synchrotron, ESRF)^18^ using protocol modified from Walsh et al.^13^ and Brunet et al.^19^ Details in supplementary materials.

### Data processing and Segmentation

Data reconstruction followed HiP-CT processing protocol as described by Brunet et al^19^. Manual segmentation was performed using Amira 2023.2 software. Figure 1 demonstrates the features of the AVCS as shown by HiP-CT. Details in supplementary materials.

**Figure 1:**
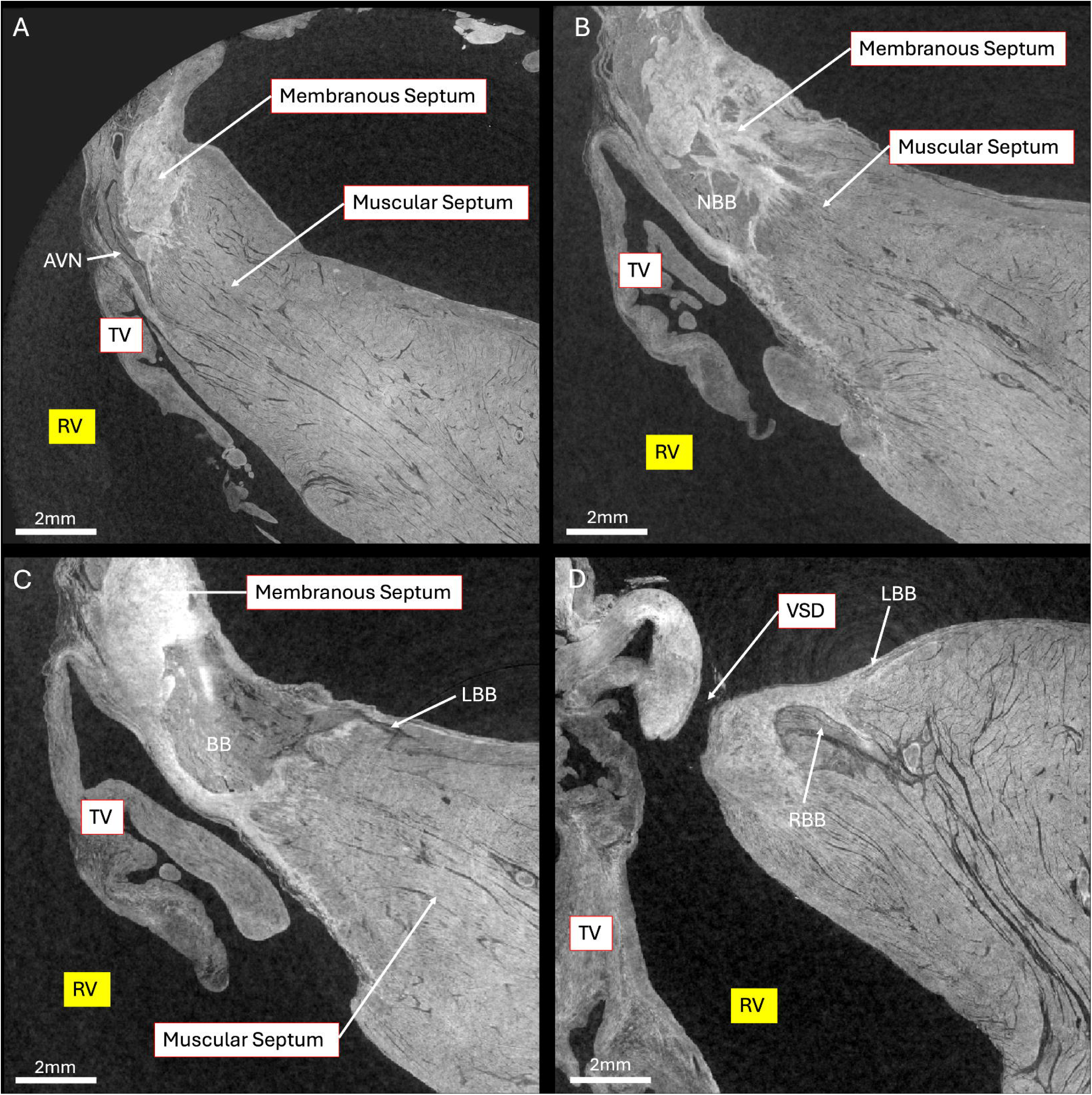
HiP-CT images of a ToF specimen with PM VSD and pulmonary stenosis in long-axis view. A – first point where the non-branching bundle (NBB) is completely ensheathed in fibrous tissue. B – left bundle branch (LBB) appears from branching bundle (BB). C – bifurcation of the right (RBB) and left bundle branches. RV – right ventricle, TV – tricuspid valve.

### Virtual Reality

Whole heart and conduction segmentations were exported separately from Amira (v2023.2) in .obj (wavefront) format and downsampled using a custom Python script to minimize mesh complexity and reach a size below 200GB (Code Availability). Exploration, visualization and manipulation of 3D models was performed in VheaRts, an extended reality (XR) application developed in Unity3D using OpenXR framework, currently used with Meta Quest devices^20,21^.

### Morphometric characteristics of the non-branching bundle and right bundle branch

NBB depth and relationship to the membranous septum remnant were quantified using Neuroglancer^22^. RBB depth and cross-sectional area through its course were quantified using an in-house Python pipeline (see Code Availability). Control and ToF samples were compared using independent t-tests.

## Results

### Hierarchical imaging

In whole-organ scans, full visualization of the AVCS was achieved in 4/7 control and 2/11 ToF specimens. The RBB was not distinguishable in the remaining datasets. In zoom scans, the AVN, NBB, BB, left bundle branch (LBB), and RBB origin could be visualized in nine ToF specimens (Supplementary table S2).

### The atrioventricular conduction system in the normal heart

The AVN was positioned at the apex of the triangle of Koch. The NBB coursed superiorly and centrally on the crest of the interventricular septum (IVS) and was aligned with the anteroseptal commissure (ASC). The LBB originated first, and the RBB arose distal to and within 2mm of the tricuspid valve (TV) annulus. The RBB quickly penetrated the myocardium and remained closely associated with the inferior limb of the septomarginal trabeculation (SMT) as it extended inferiorly towards the medial papillary muscle (MPM) or cords. The LBB exhibited variable morphology; sheet-like in some, and narrower in others. Overall, the LBB was relatively compact and positioned subendocardially (Figure 2).

**Figure 2:**
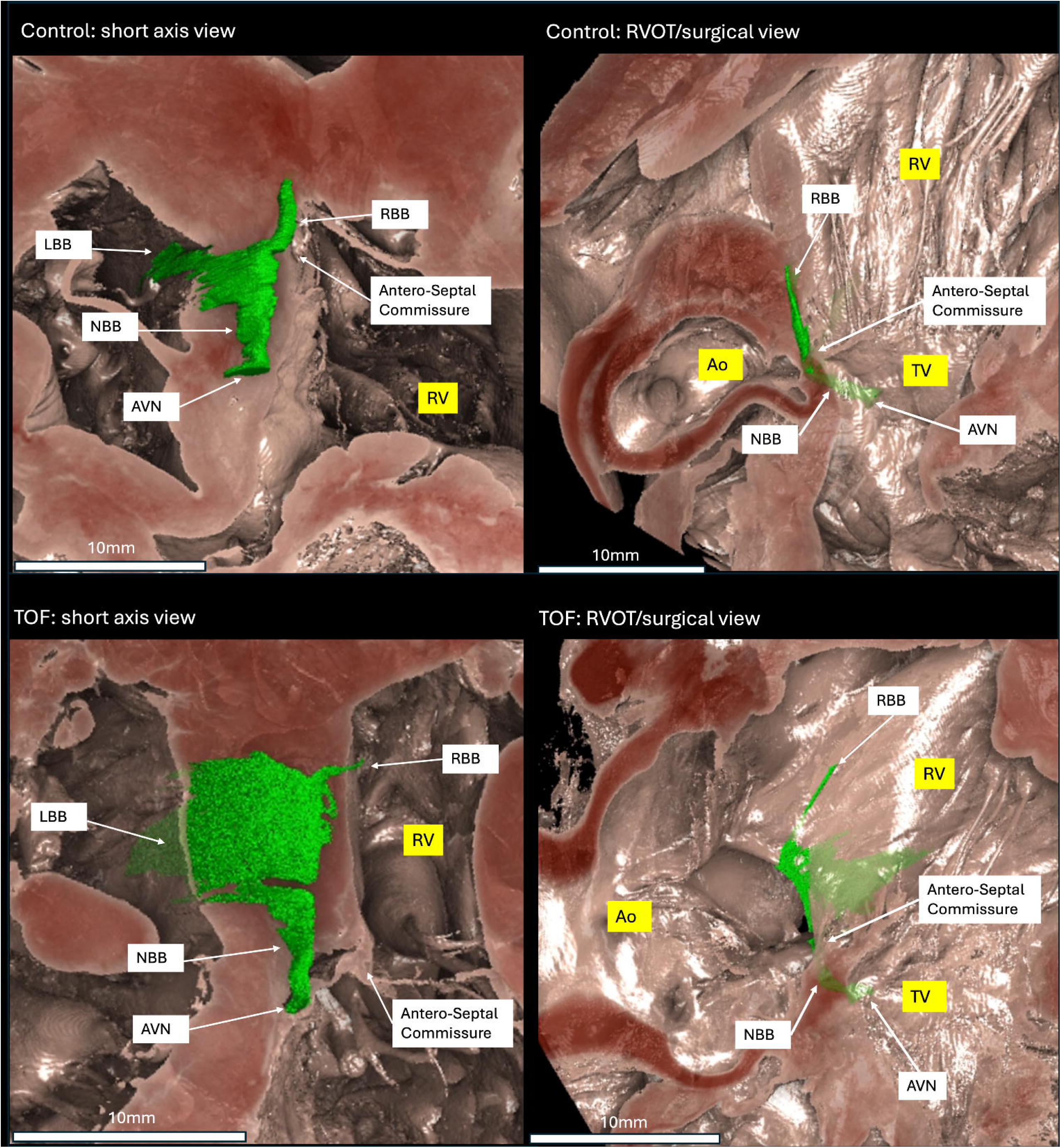
The conduction system in a control specimen in short axis and RVOT views and in a ToF specimen. AVN – atrioventricular node, NBB – non-branching bundle, RBB – right bundle branch, LBB – left bundle branch, TV – tricuspid valve, RV – right ventricle, Ao-aorta, medial PM – medial papillary muscle.

### The conduction system in Tetralogy of Fallot

The ToF AVCS exhibited significant heterogeneity. The AVN remained at the apex of the triangle of Koch, reflecting correct alignment of the atrial and ventricular septa. The NBB deviated to the left side of the IVS in four specimens (T7, T8, T10, T11), to the right in one (T1), and in four was central (T3-T6). In two (T2, T9), the NBB region was fragmented. In perimembranous VSD samples, the NBB was positioned between the membranous and muscular septa. The NBB originated on the atrial side of the TV annulus in six specimens (T2-4, T7, T9, T11) aligned with the ASC in two (T1, T8) and on the ventricular side in three (T5, T6, T10). In the inferior corner of the PM VSD (the region of mitral–tricuspid continuity), the NBB was less than 1 mm below the endocardial surface (0.12 ± 0.09 mm [median 0.09; range 0.005–0.30]). At the TV hinge line, conduction tissue lay deeper: average depth of 0.49 ± 0.30 mm [median 0.40; range 0.20–1.27]. Dead-end tracts were found in six specimens (T1-3, T6, T9, T10), resembling a ‘saddle-shaped’ extension, following the VSD contour. One large fasciculo-ventricular tract was found (T5).

In four specimens (T1, T2, T4, T6), the RBB emerged closely associated with the ASC; in the rest, it branched later without a distinctive anatomical landmark. The RBB course was associated with the MPM in 6 of 11 specimens (T1-3, T7, T8, T11) and the remainder coursed inferior to the caudal limb of SMT, unrelated to the MPM. The RBB was broad in two samples(T6, T7), draping extensively either over the right side of the septal crest or fanning bilaterally over the septal crest; while in another, it was crescent-shaped (T5). Representations of the ToF AVCS are found in Figures 3 and 4. 3D renderings are found in Figure 5, Video 1, and Supplementary Figure 1.

**Figure 3:**
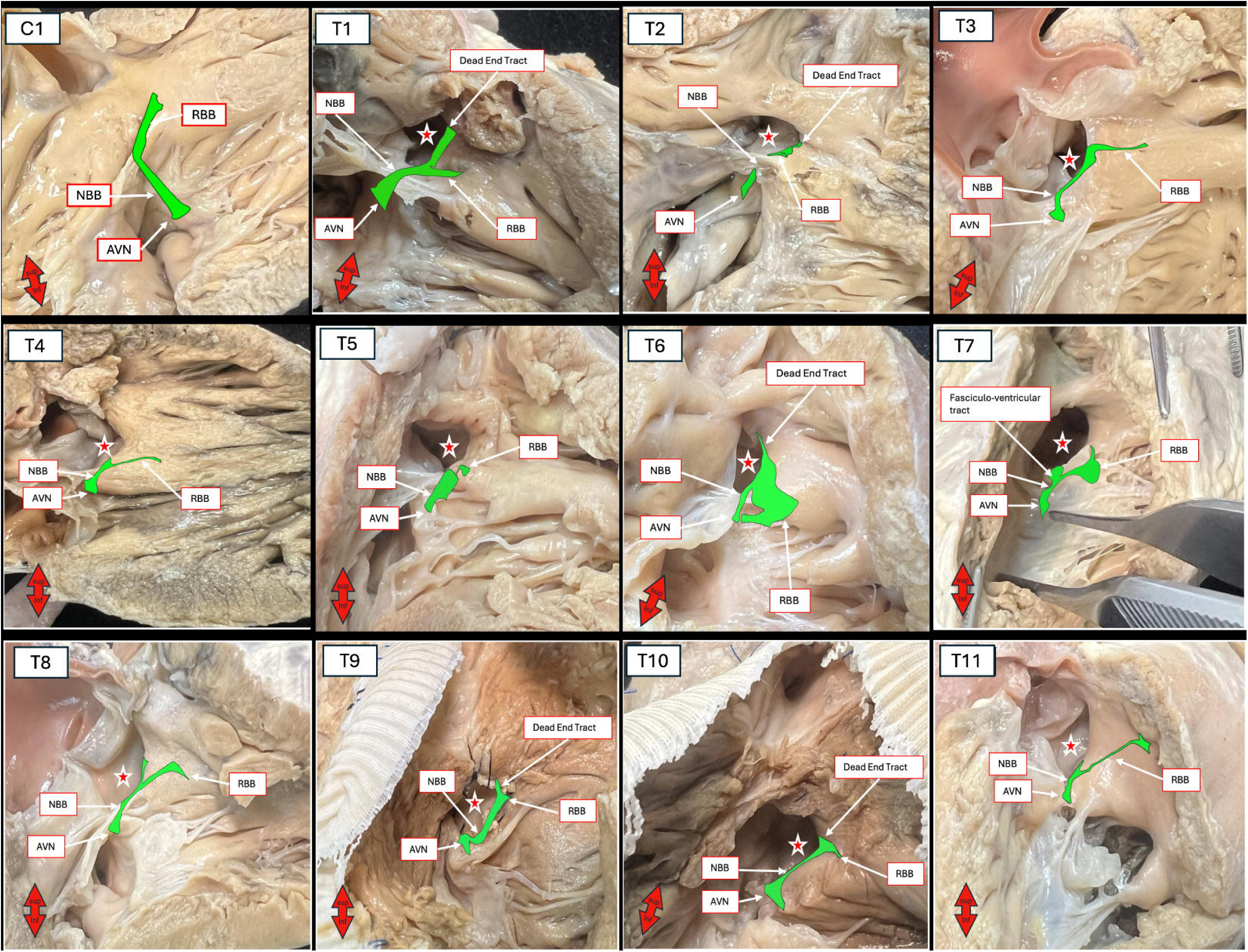
atrio-ventricular conduction system represented on a control specimen and ToF specimens. AVN – atrioventricular node, NBB – non branching bundle, RBB – right bundle branch, LBB – left bundle branch, red star – ventricular septal defect. Compass demonstrates the image orientation – inferior pointing towards diaphragmatic surface of the heart.

**Figure 4:**
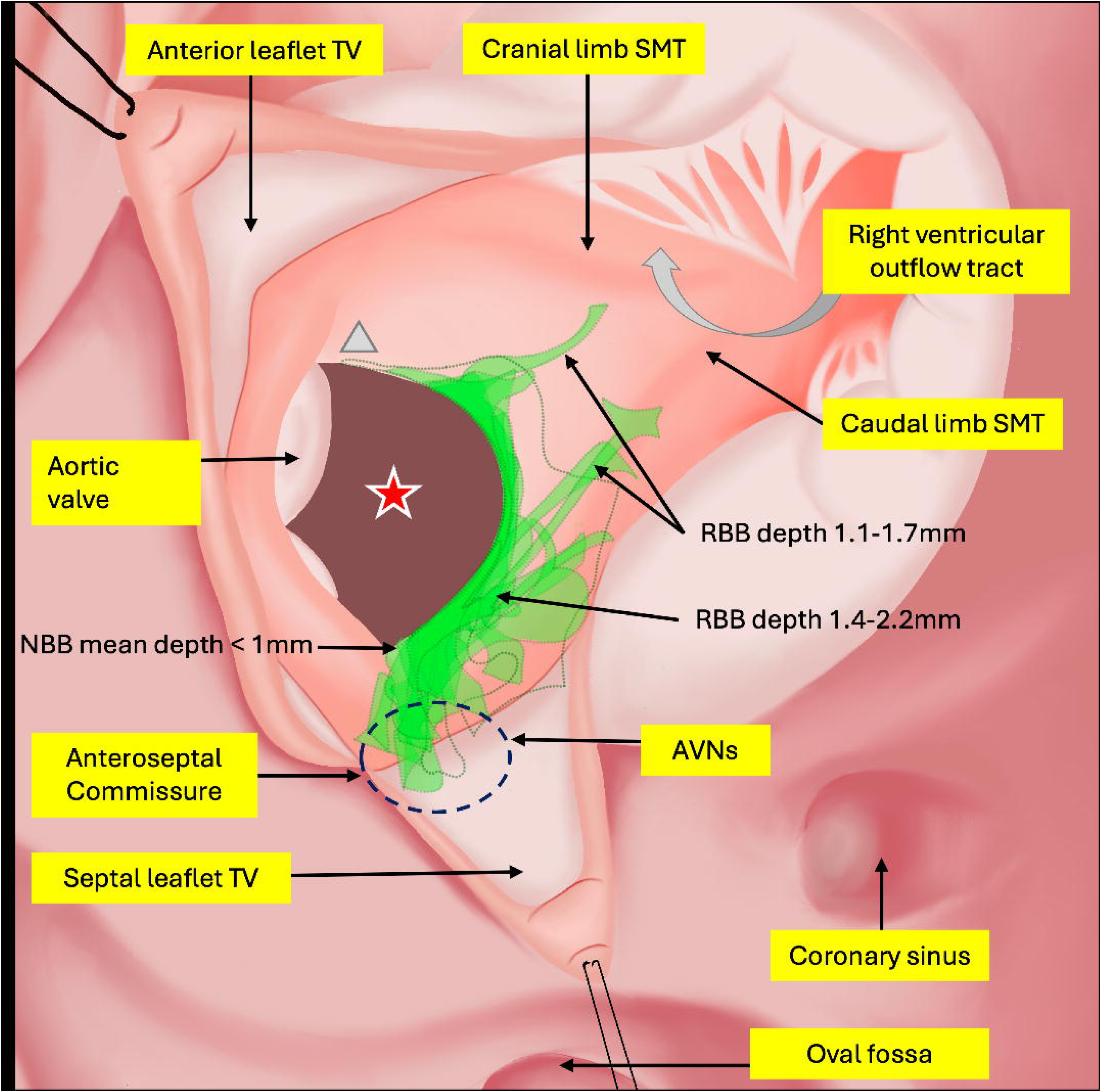
Representation of all ToF right bundle branches in this study from surgical view with the depth at relevant locations noted. NBB – non-branching bundle, RBB – right bundle branch, RVOT – right ventricular outflow tract, SMT – Septomarginal trabeculation, TV – tricuspid valve, red star – VSD, grey triangle – conduction free zone.

**Figure 5.**
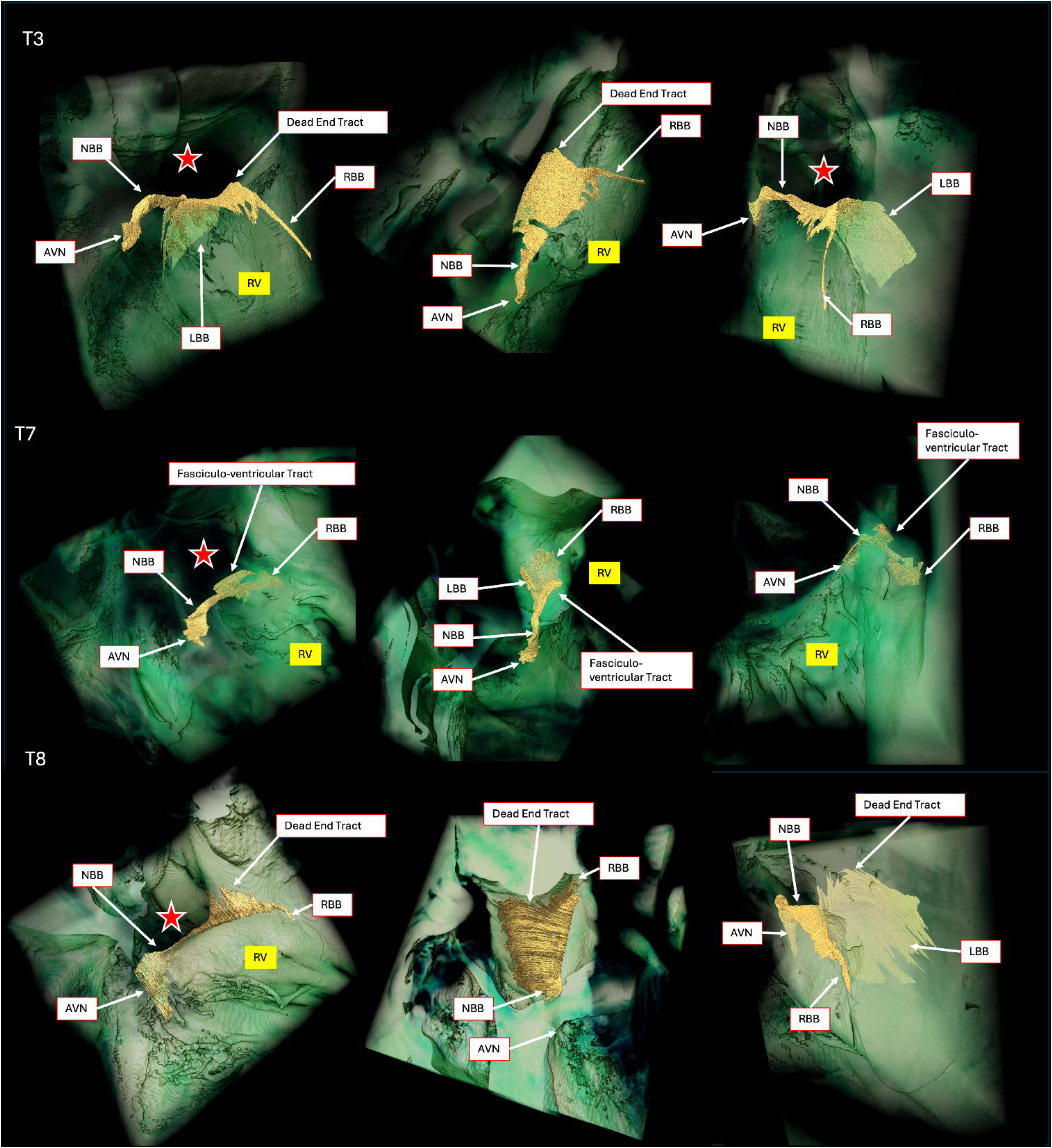
3D representations of the atrioventricular conduction system in three ToF samples. AVN – atrioventricular node, NBB – non branching bundle, RBB – right bundle branch, LBB – left bundle branch, RV – right ventricle, red star – ventricular septal defect.

### Association of conduction tissue with the membranous remnant of the interventricular septum

In PM VSDs, the mean height of the membranous septum remnant was 0.9 ± 0.19mm. In three (T4, T5, T7), conduction tissue penetrated the muscular septum below the level of the VSD, leaving the membranous septum and surrounding regions free of conduction tissue. In the remaining samples and in the muscular VSD case, conduction tissue ran between the membranous and muscular septa.

### Morphometric characteristics of the right bundle branch

In normal samples, the RBB average depth throughout its course ranged between 0.6-1.2 mm. The RBB became more superficial and then penetrated the myocardium approximately 1 mm from its origin. In ToF specimens, the RBB was deeper, ranging from 1-2.2 mm, but showed variability throughout its course (Fig 4).

Overall, ToF hearts demonstrated significantly smaller RBB cross-sectional areas compared with controls (0.12 ± 0.13 mm² [median 0.063; range 0.0001–0.36] vs 0.50 ± 0.17 mm² [median 0.50; range 0.21–0.72]; p = 0.0005). When indexed to ventricular length, similar results were observed (3.0 ± 3.1 ×10U³ mm²/mm [median 1.9; range 0.003–8.8 ×10U³] vs 16.8 ± 6.2 ×10U³ mm²/mm [median 17.5; range 5.8–24.1 ×10U³]; p = 0.0006) (Figure 6).

**Figure 6.**
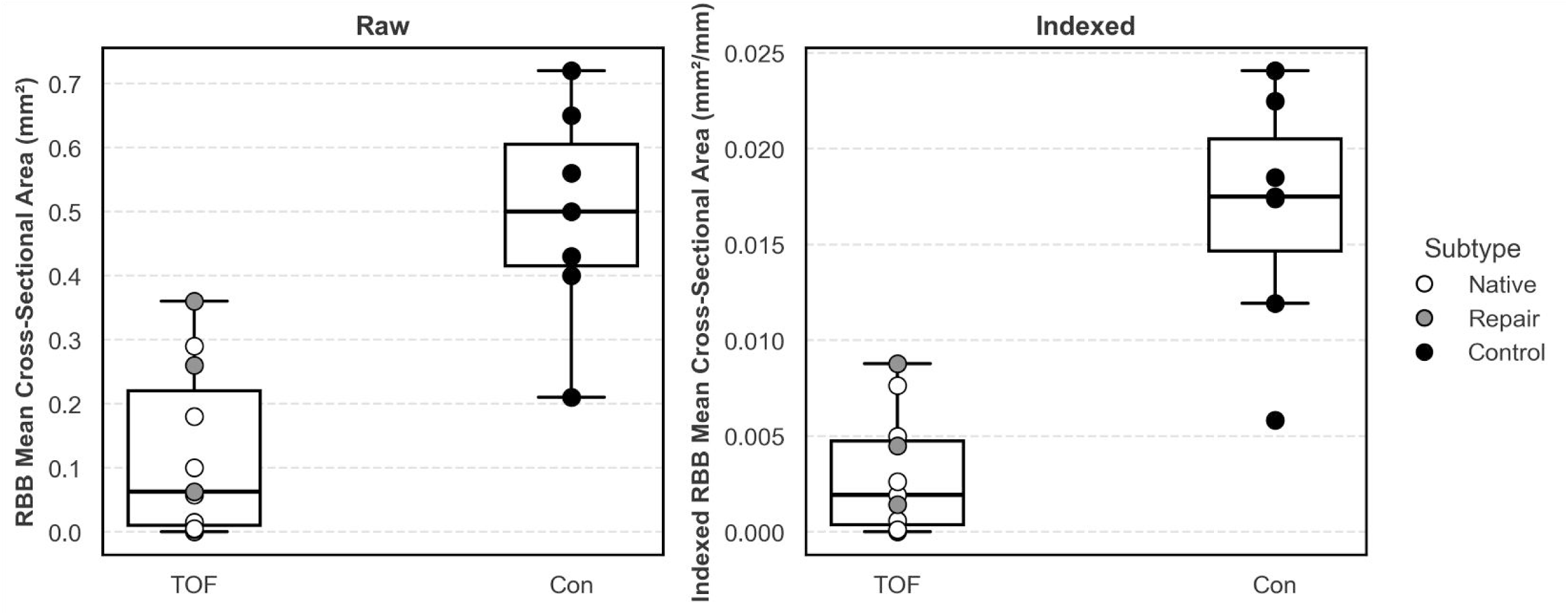
Right bundle branch mean cross-sectional area in ToF and Control hearts in raw and indexed (against ventricular length) datasets.

### Virtual Reality as a clinically relevant output

The HiP-CT to VR pipeline was successfully implemented, as proof of concept, in all trialled datasets (10 ToF, 1 Control). VR allowed detailed exploration of the AVCS and reslicing of the data in any plane as a supplementary aid to understand the spatial relationship of the AVCS to key neighboring structures (MPM, ASC, aortic valve), and to trace complex pathways in 3D (supplementary video).

## Discussion

This first HiP-CT application to a pediatric congenital heart disease case series provides new 3D data concerning the ToF AVCS. HiP-CT delivers near-histological resolution while preserving whole-organ spatial context and maintaining manageable data size, enabling 3D reconstruction and visualization of the entire heart and AVCS, with the capability for segmentation and examination in any plane. Such HiP-CT data has a potential use in hands-on surgical training (3D prints and XR), as a training set for machine learning, and for complex electrophysiologic simulations. Conventional histology relies on destructive methods and 3D reconstruction based on limited 2D data, risking anatomical misinterpretation or missing significant features.

Our findings corroborate previous studies regarding the anatomical position of the ToF NBB. The NBB is closely associated with the ASC and typically lies centrally or towards the left side of the IVS. In ToF with perimembranous VSDs, the NBB lies between the membranous and muscular septa^7^. The NBB is superficial and therefore at risk during transatrial TV leaflet retraction or VSD closure: the region of fibrous continuity between the atrio-ventricular valves at the posterior-inferior VSD margin is a recognized ‘danger’ zone in PM defects^7,8,23^. We show that conduction tissue lies within 1-3mm of the membranous septum remnant. Suturing here therefore requires caution.

The left side of the septal crest was entirely draped by conduction tissue in all ToF specimens. In some, the BB draped into the RV below the crest. The RBB’s variable origin, anywhere from the TV annulus to the VSD nadir, highlights the need for caution in this region. Using the intersection of the VSD with the aortic leaflets as a reference point, no branching was observed between this point and 2mm beyond the start of the curvature of the VSD nadir. No correlation was found between RBB origin location and its subsequent course in ToF, further complicated by the inconsistent association of the RBB with the MPM. RBB depth from the endocardial surface at origin ranged from 0.25 mm to 2.5 mm, becoming shallower within 1–3 mm. Although variable depth endangers the ToF RBB during surgery, its narrow nature could be protective.

Suturing 3-4 mm below the IVS crest, between the upwards curvature and nadir, poses moderate risk. A balance between suturing inferiorly and maintaining stability of the VSD patch is important. Suturing tangentially forehand or backhand parallel to the IVS crest could reduce risk.

The ToF RBB was less traceable than in controls, and in two samples, the RBB remained invisible even at 2.203 μm/voxel, possibly because the RBB branches more proximally into thin, sheet-like structures which are challenging to trace even at the highest resolution HiP-CT currently available.

The ToF dead-end tracts were continuations of the BB, following the VSD curvature towards the aortic root. Limited descriptions of similar tracts in literature are primarily in structurally normal hearts and in one ToF case^24,25^. In one sample, an atypical, sheet-like fasciculo-ventricular tract was observed running superiorly. Fasciculo-ventricular connections, ubiquitous in normal and congenitally malformed hearts, can create re-entry circuits, independent of AVN input. These can be insulated and bypass normal conduction pathways, enabling atypical forms of ventricular pre-excitation. The persistence of abnormal tracts in ToF, accurately visualized by HiP-CT, appears more common than previously described and could represent an under-recognized substrate for spontaneous ventricular arrhythmias and unsuccessful ablation outcomes^26,27^.

Our observations of a less well-defined AVCS in ToF and additional anomalies, suggests abnormal differentiation during development, potentially providing a larger substrate for both pre- and post-operative arrhythmias, driven by enhanced automaticity or triggered activity within the atrioventricular junction (e.g JET), or for ectopic impulse generation and re-entry. In many respects, the ToF AVCS resembles early stages of cardiac development, for example, at Carnegies stage 16/17 in the human embryo, before septal closure and remodelling of conduction tissue in relation to the IVS crest^28–30^. These anomalies may reflect altered differentiation/remodelling of cardiomyocytes into conduction tissue, driven by key transcription factors^31,32^. Collaborative work is currently underway to explore such mechanisms, combining spatial transcriptomic data with 3D volumes obtained through HiP-CT.

This study demonstrates proof of concept for a complete pathway integrating HiP-CT segmentations into VR. Although 3D reconstructions can be produced in specialized software, interpretation on 2D screens remains challenging. Here, VR was shown to be a useful aid for visualizing and manipulating complex cardiac anatomy, allowing the AVCS to be examined in relation to surrounding heart structures. In future, it may be possible in VR to train surgeons to avoid the AVCS in ToF through surgical or patch planning simulations. Further development and validation of the HiP-CT to VR pipeline is warranted to establish its clinical and educational utility.

This study shows that heterogeneity of the ToF AVCS makes its course unpredictable, highlighting the value of intraoperative conduction mapping. Successful localization of the NBB using high-density multi-electrode catheters has been achieved and shows promise in reducing post-operative atrioventricular block^33,34^. Expanding intraoperative mapping to detect the RBB could further reduce surgical risk.

### Limitations

Phase-contrast tomography at synchrotrons remains an expensive and limited-access technique, restricted to ex-vivo imaging due to the need for absorption normalization, auto-absorption protocols, and the very high radiation dose. Lab-based systems under development may broaden access. The relatively small and diverse sample set limited further sub-analyses. Despite careful selection of intact specimens, handling over time occasionally caused unexpected fractures and fragmentation of conduction tissue not visible endocardially. The 2.203 μm/voxel ‘zooms’ did not always resolve distal conduction tissue. Higher resolution might amplify noise rather than signal. A major bottleneck is the segmentation process, which is time-consuming and requires expert morphologists. Development of machine learning models for automatic segmentation is currently underway in group, using ground truth data from this study.

## Conclusion

Using HiP-CT imaging of 18 whole hearts (11 ToF, 7 controls), coupled with custom morphometric pipelines and virtual-reality visualization, we demonstrate significant heterogeneity of the ToF conduction system, including variable right bundle branch origin, depth, and hypoplasia; draped and immature conduction morphology; the presence of dead-end tracts and fasciculo-ventricular connections; and define precise regions of surgical vulnerability around the ventricular septal defect. These findings redefine the anatomic substrate in ToF and have direct relevance for defining preoperative risk, informing surgical techniques, and demonstrating the importance of intraoperative conduction mapping strategies.

## Supporting information

Details in supplementary materials.

## Code Availability

All analysis and utility scripts used in this study are openly available at the public repository: https://github.com/VaishnaviSabari/conduction-tools.git.

## Data Availability

DOIs for data used in this paper are available on request and are hosted on the ESRF data repository at https://human-organ-atlas.esrf.fr/.

## Abbreviations

ASC: anteroseptal commissure
AVCS: atrioventricular conduction system
AVN: atrioventricular node
BB: Branching Bundle
BCH: Birmingham Children’s Hospital
ESRF: European Synchrotron Radiation Facility
HiP-CT: Hierarchical Phase-Contrast Tomography
IVS: interventricular septum
LBB: Left bundle branch
MPM: medial papillary muscle
NBB: non-branching bundle
PM: perimembranous
RBB: Right bundle branch
ToF: Tetralogy of Fallot
TV: Tricuspid Valve
VSD: ventricular septal defect
JET: Junctional Ectopic Tachycardia
ZCR: Zayed Centre for Research

---

Supplementary Figure 1: 3D renderings of conduction system segmentations in one control, and all ToF specimens: AVN – atrioventricular node, NBB – non-branching bundle, RBB – right bundle branch, LBB – left bundle branch, RV – right ventricle, star - VSD.

