## Supplementary material for "The Heterogeneous Nature of Atrioventricular Conduction Tissues in Tetralogy of Fallot demonstrated by Hierarchical Phase-contrast Tomography – redefining the anatomic substrate": Details in supplementary materials.

### Sample Preparation and Data Acquisition

Each heart was mounted in a container filled with 4% neutral buffered formalin and agar to stabilize the sample and prevent motion during acquisition. To minimize the formation of bubbles caused by radiation exposure, specimens underwent in-line degassing prior to imaging. The containers were then hermetically sealed until scanning started.

A filtered parallel polychromatic X-ray beam was used with a 178m source-to-sample distance and propagation distances of 10m (overview) and 1m/1.6m (local zoom)^1^. X-rays were converted to visible light using LuAG:Ce scintillators (Crytur, Czechia) of 2000 μm (overview) or 50-100 μm (zoom) thickness, demagnified by Dzoom optic (overview) or magnified by a fixed x2 optic (zooms), and detected by an Iris 15 camera (Teledyne Photometrics, United States). Specimens were scanned entirely at voxel sizes of 19.19 μm (ToF with lungs), 10 μm (ToF without lungs, and ZCR normal specimens), and 7.032 μm (BCH normal controls), followed by local ‘zoom’ scans in regions of interest at 2.2 μm voxel resolution in all ToF samples and three control samples. The difference in overview pixel sizes is related to the size of the specimen. For each acquisition, a scan of a sealed container filled with 4% agar/formalin with the same parameters to serve as a beam reference for flat-field correction and correct low-frequency artefacts. Full scan parameters are in supplementary table S1.

### Data processing and analysis

Data reconstruction was performed using tomography processing software (night-rail^2^, Nabu^3^) developed at ESRF, following the HiP-CT processing protocol as described by Brunet et al^1^. The night-rail framework integrates normalization, reference pairing, vertical concatenation (or helical) and ring artefact correction using an updated version of the Lyckegaard algorithm^4^. A 2D unsharp mask was applied to enhance contrast. Processing and visualization of the large datasets required high-performance computing resources^1^.

Images were preprocessed by optimizing image brightness and contrast. 16-bit TIFF stacks were converted to 8-bit using ImageJ v1.54g^5^ to reduce size and processing time. Conduction tissue was identified for segmentation following the rules of Aschoff and Monckeberg and promoted by Anderson et al^6^: conduction myocytes were histologically discrete from working myocardium, serially traceable section to section, and insulated by a fibrous sheath. The non-branching bundle (NBB) (of His) was first located in a four-chamber plane, where it is characteristically surrounded by connective tissue, connecting inferiorly to the atrioventricular node (AVN) and superiorly to the branching bundles. In Amira (2023.2) an unsharp mark and thresholding were applied to enhance tissue definition. Conduction tissue was manually segmented every 5^th^ slice and interpolated using an in-house ImageJ macro (see Code Availability). 3D renderings were generated in Amira 2023.2 and VGStudioMax 2024.2.

Table S1: Scanning parameters for each HiP-CT scan

| Sample ID | Voxel size (um) | Propagation distance (m) | Attenuator | Mean energy (keV) | Lateral Field of View (mm) | Projections | Exposure Time (ms) | Accumulation | Acquisition Mode | Scan Time (s/scan) | Total number of scans | Vertical field of view/scan (mm) | Vertical translation |
| --- | --- | --- | --- | --- | --- | --- | --- | --- | --- | --- | --- | --- | --- |
| **125** | 9.595 | 10 | Sapphire 5mm Mo 0.21mm | 96 | 81.14 | 12000 | 20 | 1 | Half-acquisition | 265 | 15 | 8.16 | 7 |
|  | 2.21 | 1 | Sapphire 5mm, sapphire 5mm, Ag 0.1mm | 86.6 | 13.44 | 15000 | 20 | 1 | Half-acquisition | 107 | 17 | 2.94 | 1.5 |
| **173** | 9.595 | 10 | Sapphire 5mm Mo 0.21mm | 96 | 81.14 | 12000 | 20 | 1 | Half-acquisition | 265 | 14 | 8.16 | 7 |
|  | 2.203 | 1.6 | Sapphire 5mm, Ag 0.2mm | 89.6 | 19.95 | 15000 | 35 | 1 | Half-acquisition | 616 | 5 | 6.52 | 4 |
| **185** | 9.595 | 10 | Sapphire 5mm, Mo 0.21mm | 96 | 81.14 | 12000 | 20 | 1 | Half-acquisition | 265 | 15 | 8.16 | 7 |
|  | 2.203 | 1.6 | Sapphire 5mm, Ag 0.2mm | 89.6 | 19.95 | 15000 | 35 | 1 | Half-acquisition | 618 | 1 | 6.62 | 4 |
| **450** | 9.595 | 10 | Sapphire 5mm Mo 0.21mm | 96 | 81.14 | 12000 | 20 | 1 | Half-acquisition | 265 | 18 | 8.16 | 7 |
|  | 2.203 | 1.6 | Sapphire 5mm, Ag 0.2mm | 89.6 | 19.95 | 15000 | 35 | 1 | Half-acquisition | 616 | 6 | 6..52 | 4 |
| **727** | 9.595 | 10 | Sapphire 5mm, Mo 0.21mm | 96 | 81.14 | 12000 | 20 | 1 | Half-acquisition | 265 | 16 | 8.16 | 7 |
|  | 2.203 | 1.6 | Sapphire 5mm, Ag 0.2mm | 89.6 | 19.95 | 15000 | 35 | 1 | Half-acquisition | 618 | 1 | 6.52 | 4 |
| **769** | 20.01 | 10 | Sapphire 10mm, Ag 0.2mm, C 25mm | 108 | 117.9 | 12000 | 10 | 4 | Quarter-acquisition | 530 | 15 x 2 | 8.00 | 7 |
|  | 2.203 | 1.6 | Sapphire 5mm, Ag 0.2mm | 86.2 | 19.95 | 15000 | 40 | 1 | Half-acquisition | 704 | 6 | 6.52 | 4 |
| **1770** | 9.595 | 10 | Sapphire 5mm Mo 0.21mm | 96 | 81.14 | 12000 | 20 | 1 | Half-acquisition | 265 | 15 | 8.16 | 7 |
|  | 2.203 | 1 | Sapphire 10mm, Ag 0.1mm | 86.6 | 19.95 | 15000 | 20 | 1 | Half- acquisition | 4030 | 10 | 3.09 | 1.5 |
| **1771** | 9.595 | 10 | Sapphire 5mm Mo 0.21mm | 96 | 81.14 | 12000 | 20 | 1 | Half-acquisition | 265 | 15 | 8.16 | 7 |
|  | 2.203 | 1.6 | Sapphire 5mm, Ag 0.2mm | 89.6 | 19.95 | 15000 | 35 | 1 | Half-acquisition | 616 | 6 | 6.52 | 4 |
| **1840** | 19.19 | 12 | Sapphire 5mm Mo 3.75mm SiO2 bars 25mm | 155 | 137 | 12000 | 13 | 3 | Quarter-acquisition | 516 | 21 x 2 | 8.16 | 7 |
|  | 2.203 | 1.6 | Sapphire 10mm, Ag 0.1mm | 86.2 | 19.95 | 15000 | 40 | 1 | Half-acquisition | 704 | 12 | 6.52 | 4 |
| **2010** | 19.19 | 12 | Sapphire 5mm Mo 3.75mm SiO2 bars 25mm | 155 | 137 | 12000 | 13 | 3 | Quarter-acquisition | 516 | 16 x 2 | 8.06 | 7 |
|  | 2.203 | 1.6 | Sapphire 10mm, Ag 0.1mm | 86.2 | 19.95 | 15000 | 40 | 1 | Half-acquisition | 699 | 9 | 6.52 | 4 |
| **2577** | 10.005 | 10 | Sapphire 10mm Ag 0.2mm C 20mm | 105 | 90.6 | 12000 | 20 | 1 | Half-acquisition | 265 | 16 | 8.00 | 7 |
|  | 2.203 | 1.6 | Sapphire 5mm, Ag 0.2mm | 89.6 | 19.95 | 15000 | 35 | 1 | Half-acquisition | 617 | 1 | 6.52 | 4 |
| **2359** | 9.595 | 10 | Sapphire 5mm Mo 0.21mm | 96 | 81.14 | 12000 | 20 | 1 | Half-acquisition | 265 | 14 | 8.16 | 7 |
| **2337** | 10.005 | 10 | Sapphire 10mm Ag 0.2mm C 20mm | 108 | 90.6 | 15000 | 20 | 3 | Half-acquisition | 1026 | 15 | 8.9 | 7 |
| **2625** | 7.032 | 7 | Sapphire 10mm, Ag 0.2mm, glassy carbon 30mm | 105 | 66.5 | 15000 | 20 | 3 | Half-acquisition | 1023 | 11 | 8.02 | 7 |
|  | 2.203 | 2.1 | Sapphire 10, Ag 0.3mm, glassy carbon 30mm | 94.5 | 19.95 | 15000 | 50 | 1 | Half-acquisition | 872 | 4 | 6.52 | 4 |
| **2627** | 7.032 | 7 | Sapphire 10mm, Ag 0.2mm, glassy carbon 30mm | 105 | 66.5 | 15000 | 20 | 3 | Half-acquisition | 1027 | 12 | 8.02 | 7 |
|  | 2.203 | 2.1 | Sapphire 10mm, Ag 0.2mm, glassy carbon 30mm | 94.5 | 19.95 | 15000 | 50 | 1 | Half-acquisition | 871 | 1 | 6.52 | 4 |
| **2628** | 7.032 | 7 | Sapphire 10, Ag 0.3mm, glassy carbon 30mm | 105 | 66.5 | 15000 | 20 | 3 | Half-acquisition | 1027 | 12 | 8.02 | 7 |
| **2629** | 7.032 | 7 | Sapphire 10mm, Ag 0.2mm, glassy carbon 30mm | 105 | 66.5 | 15000 | 20 | 3 | Half-acquisition | 1027 | 9 | 8.02 | 7 |
|  | 2.201 | 2.1 | Sapphire 10mm, Ag 0.3mm, glassy carbon 9.6mm | 94.5 | 19.95 | 15000 | 50 | 1 | Half-acquisition | 872 | 1 | 6.52 | 4 |
| **2630** | 7.032 | 7 | Sapphire 10mm, Ag 0.2mm, glassy carbon 30mm | 105 | 66.5 | 15000 | 20 | 3 | Half-acquisition | 1027 | 9 | 8.02 | 7 |

Table S2: AV conduction system structures visible in original whole-organ dataset (black ticks) and in local high-resolution zoom scans 2.21um/2.203um (red ticks).

| Sample | AVN | PB | LBB | RBB origin | RBB | Non-branching bundle |
| --- | --- | --- | --- | --- | --- | --- |
| **125** | 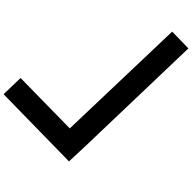 | 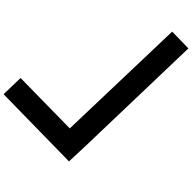 | 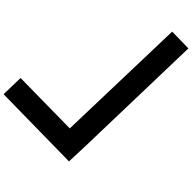 | 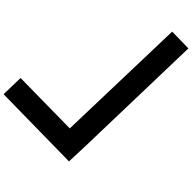 | 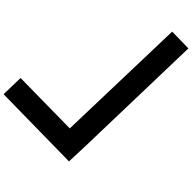 | 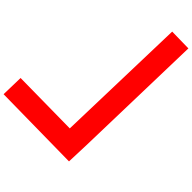 |
| **173** | 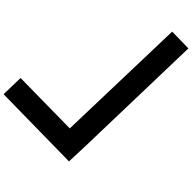 | 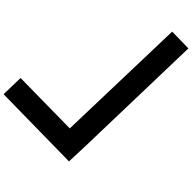 | 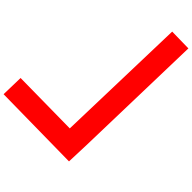 | 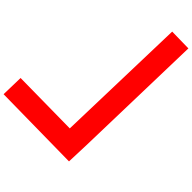 | 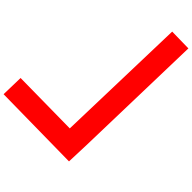 | 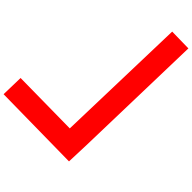 |
| **185** | 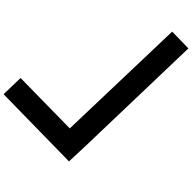 | 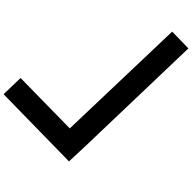 | 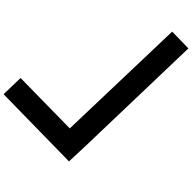 | 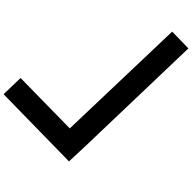 | 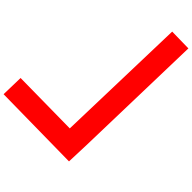 | 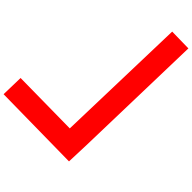 |
| **450** | 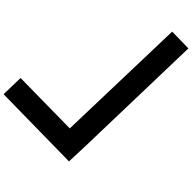 | 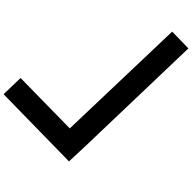 | 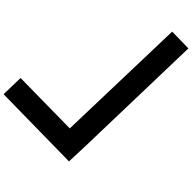 | 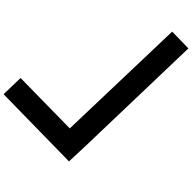 | 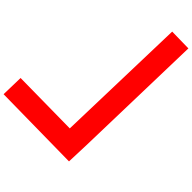 | 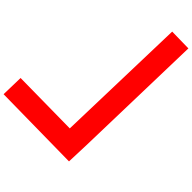 |
| **727** | 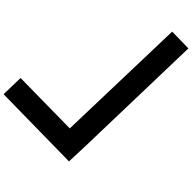 | 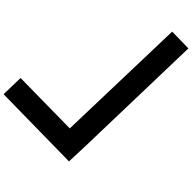 | 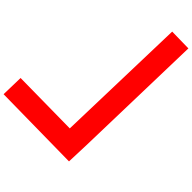 | 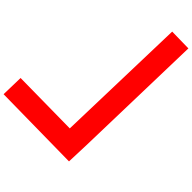 | 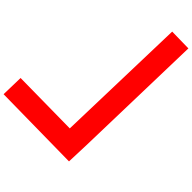 | 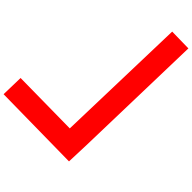 |
| **769** | 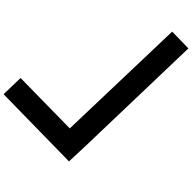 | 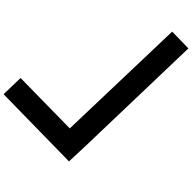 | 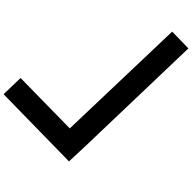 | 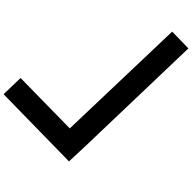 | 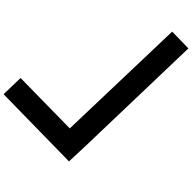 | 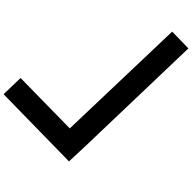 |
| **1770** | 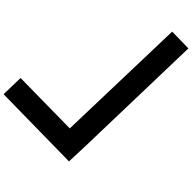 | 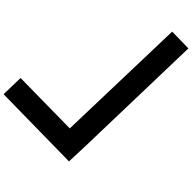 |  | 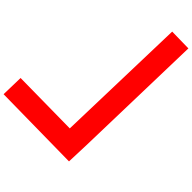 | 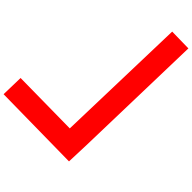 | 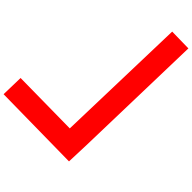 |
| **1771** | 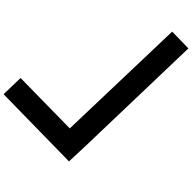 | 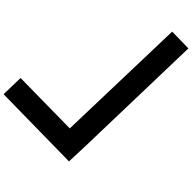 | 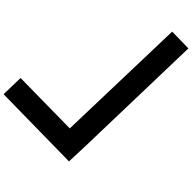 | 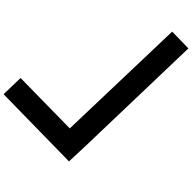 | 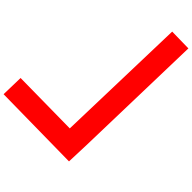 | 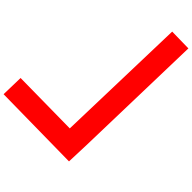 |
| **1840** | 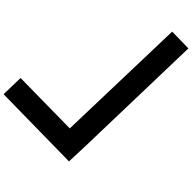 | 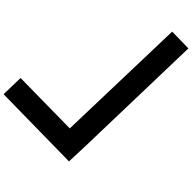 | 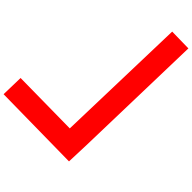 | 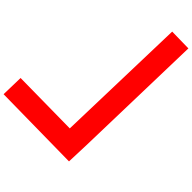 | 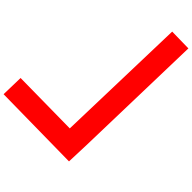 | 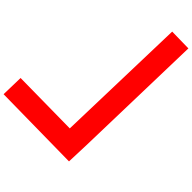 |
| **2010** | 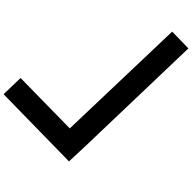 | 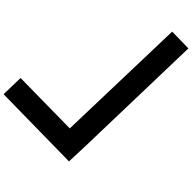 | 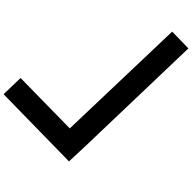 | 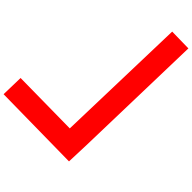 | 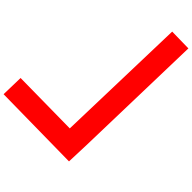 | 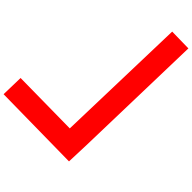 |
| **2577** | 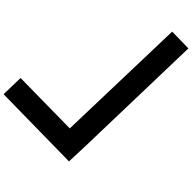 | 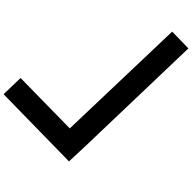 | 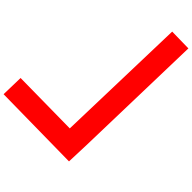 | 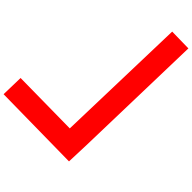 | 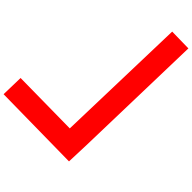 | 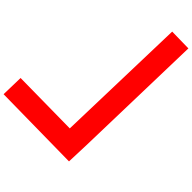 |
| **2359** | 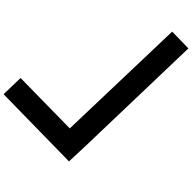 | 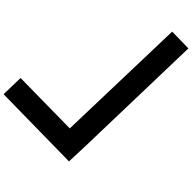 | 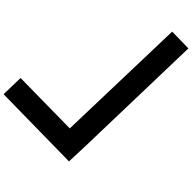 | 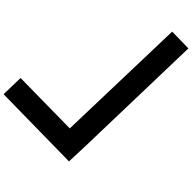 | 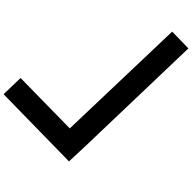 | 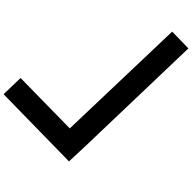 |
| **2337** | 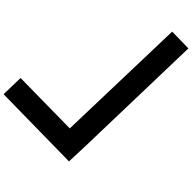 | 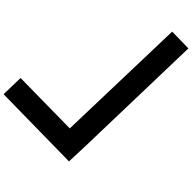 | 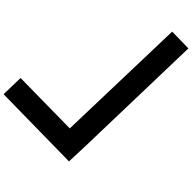 | 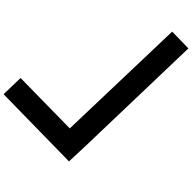 | 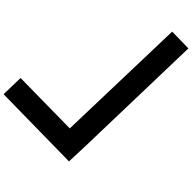 | 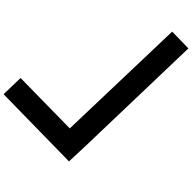 |
| **2625** | 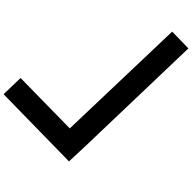 | 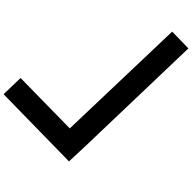 | 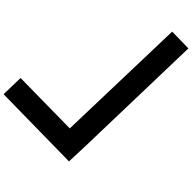 | 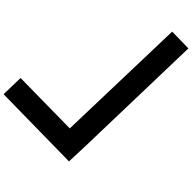 |  |  |
| **2627** | 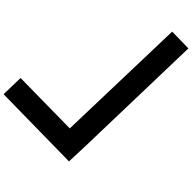 | 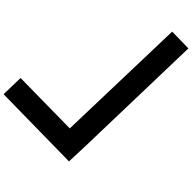 | 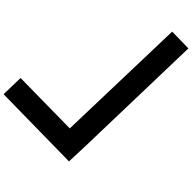 | 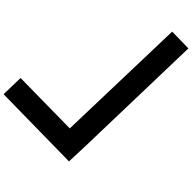 | 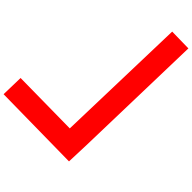 | 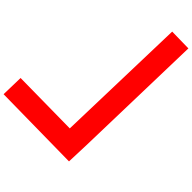 |
| **2628** | 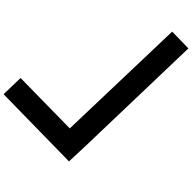 | 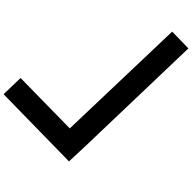 | 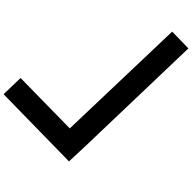 | 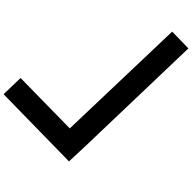 | 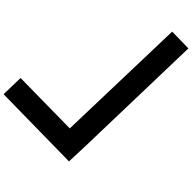 | 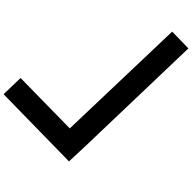 |
| **2629** | 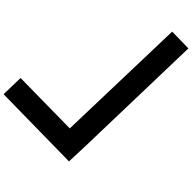 | 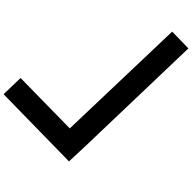 | 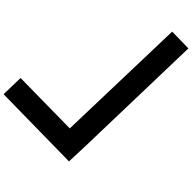 | 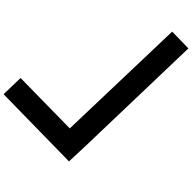 | 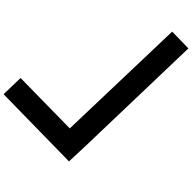 | 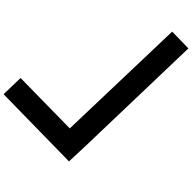 |
| **2630** | 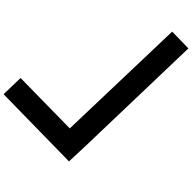 |  |  |  |  |  |
