## Supplementary figures and images for "The Heterogeneous Nature of Atrioventricular Conduction Tissues in Tetralogy of Fallot demonstrated by Hierarchical Phase-contrast Tomography – redefining the anatomic substrate"

### Supplementary Figure 1

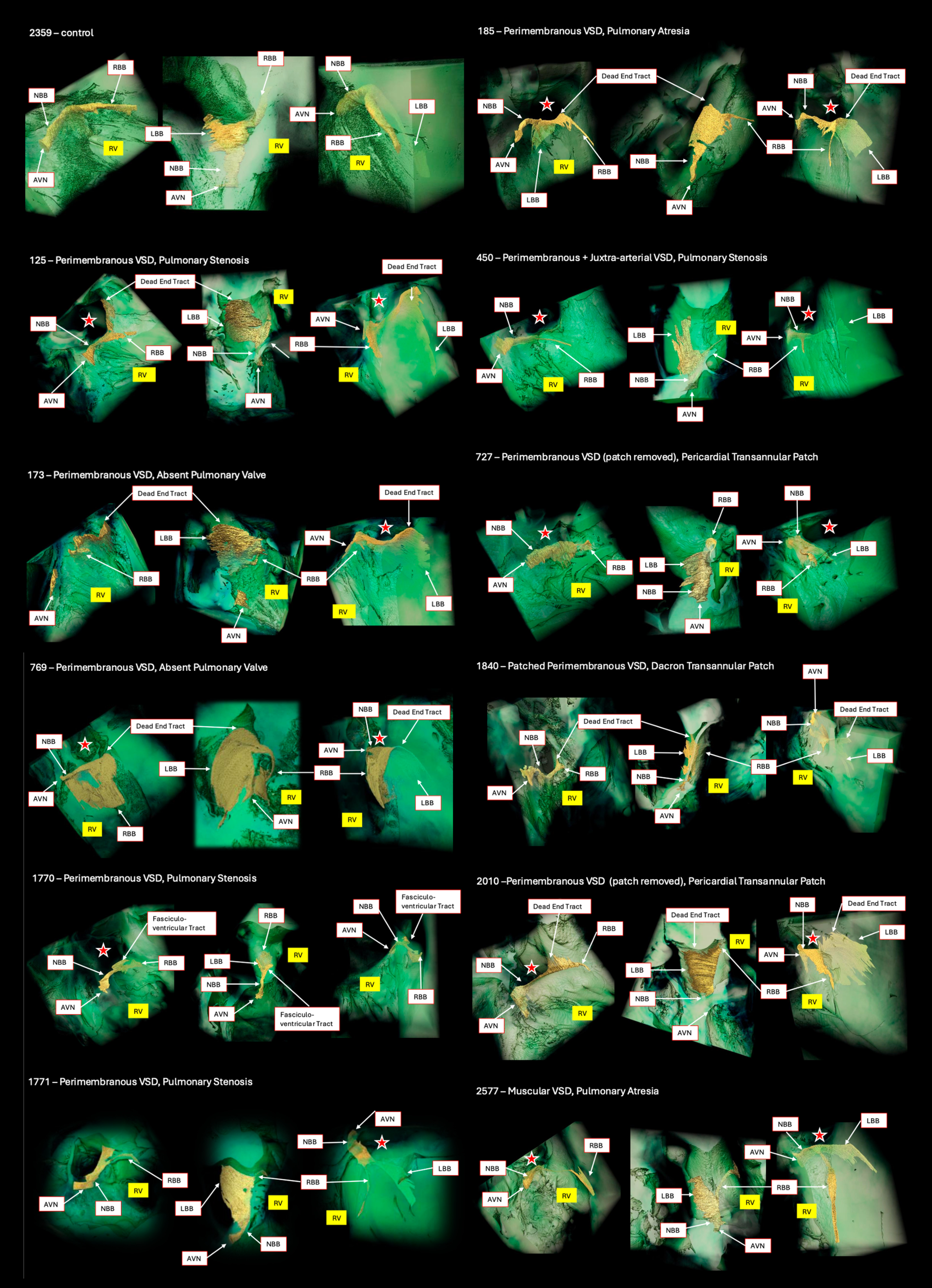
